# Pharmacological activation of constitutive androstane receptor induces female-specific modulation of hepatic metabolism

**DOI:** 10.1101/2023.04.17.537169

**Authors:** Huillet Marine, Lasserre Frédéric, Gratacap Marie-Pierre, Engelmann Beatrice, Bruse Justine, Polizzi Arnaud, Fougeray Tiffany, Martin Céline, Rives Clémence, Fougerat Anne, Naylies Claire, Lippi Yannick, Garcia Géraldine, Rousseau- Bacquie Elodie, Canlet Cécile, Debrauwer Laurent, Rolle-Kampczyk Ulrike, von Bergen Martin, Payrastre Bernard, Boutet-Robinet Elisa, Gamet-Payrastre Laurence, Guillou Hervé, Loiseau Nicolas, Ellero-Simatos Sandrine

**Affiliations:** Toxalim (Research Centre in Food Toxicology), INRAE, ENVT, INP-Purpan, UPS, Université de Toulouse, Toulouse, France; INSERM, UMR-1297 and Université Toulouse III, Institut de Maladies Métaboliques et Cardiovasculaires (I2MC), CHU-Rangueil, F-31300 Toulouse, France; Helmholtz Centre for Environmental Research, Department Molecular Systems Biology, 04318 Leipzig, Germany; CHU de Toulouse, Laboratoire d’Hématologie, Toulouse Cedex, France

**Keywords:** Sexual dimorphism, hepatic xenobiotic metabolism, lipoprotein metabolism, platelet aggregation, trimethylamine-N-oxide

## Abstract

**Background and Aims:** The constitutive androstane receptor (CAR) is a nuclear receptor able to recognize a large panel of xenobiotics leading to the modulation of the expression of its target genes involved in xenobiotic detoxication and energy metabolism. While CAR hepatic activity is thought to be higher in women than in men, its response to an acute pharmacological activation has never been investigated in both sexes.

**Methods:** Hepatic transcriptome, plasma and hepatic metabolome, have been analyzed in *Car^+/+^* and *Car^-/-^* male and female mice treated either with the CAR-specific agonist, 1,4-bis[2-(3,5-dichloropyridyloxy)]benzene (TCPOBOP), or with vehicle.

**Results:** While 90% of TCPOBOP-sensitive genes were modulated in a sex-independent way, the remaining 10% were almost exclusively impacted in female liver specifically. These female-specific CAR-sensitive genes were mainly involved in xenobiotic metabolism, inflammation and extracellular matrix organization. CAR activation also induced higher hepatic oxidative stress and hepatocyte cytolysis in females than in males. Data mining on human data confirmed that CAR activation may be involved in sexually-dimorphic drug-induced liver injury. Hepatic expression of flavin monooxygenase 3 *(Fmo3)* was almost abolished and associated with a decrease of hepatic trimethylamine-N-oxide (TMAO) concentration in TCPOBOP-treated females. In line with a possible role in the control of TMAO homeostasis, CAR activation decreased platelet hyperresponsiveness in female mice supplemented with dietary choline.

**Conclusions:** Our results demonstrate that more than 10% of CAR-sensitive genes are sex-specific and influence hepatic and systemic response such as platelet aggregation. Also, CAR activation may be an important mechanism of sexually-dimorphic drug-induced liver injury.

## Highlights

- We describe sex-independent *vs.* sex-biased CAR-dependent hepatic pathways
- CAR activation impacts liver metabolism in female mice more than in males
- CAR might play a role in women-biased drug-induced liver injury
- CAR activation inhibits TMAO synthesis, which could impact platelet aggregation

## Introduction

The liver is one of the most sexually dimorphic organs. There is increasing evidence for sexually dimorphic regulation of xenobiotic clearance, responses to drugs and drug-induced liver injury. We postulated that the mechanisms underlying such dimorphism may involve the constitutive androstane receptor (CAR, NR1I3). CAR is a liver-enriched member of the nuclear receptor superfamily that controls ligand-dependent regulation of gene expression. Upon ligand-binding, CAR translocates to the nucleus, heterodimerizes with retinoid X receptor alpha and binds the xenobiotic response element located on the DNA, upstream of the promoter sequences of its target genes. CAR was first described as a xenobiotic receptor that recognizes a wide variety of drugs, food and environmental pollutants^1, 2^. Later studies have then unveiled that CAR can also be activated by endobiotics such as bilirubin, bile acids and steroid hormones^3, 4^. Upon activation, CAR regulates the expression of critical enzymes involved in phase I, phase II and phase III xenobiotic metabolism^5, 6^, thereby playing a central role in xenobiotic detoxification and clearance. Moreover, CAR is involved in glucose and lipid homeostasis, although its exact role in hepatic metabolism remains debated^7^.

Hepatic expression of *Car* and its main target genes *Cyp2b9* and *Cyp2b10* is higher in female mice than in males^8, 9^. Moreover, treatment with nonylphenol, a moderate CAR activator, induced expression of cytochromes P450 more strongly in female mouse liver than in males^10^. Similarly, treatment with 1,4-bis[2-(3,5- dichloropyridyloxy)]benzene (TCPOBOP), the prototypical pharmacological agonist of murine CAR (mCAR)^11^, increased liver proliferation in female more than in male mice^12^. Our own previous study has shown that CAR deletion had a stronger impact on female hepatic gene expression than on that of males, however, CAR-deleted males developed spontaneous steatosis during aging while females did not^13^.Therefore, CAR is thought to impact rodent liver gene expression in a sex-dependent way, with higher CAR activity and higher sensitivity to CAR activation in females. Interestingly, in humans, the expression and activity of CYP2B6, the prototypical target gene for human CAR (hCAR), were higher in women’s liver compared to men, revealing that sexual dimorphism observed in CAR activity also seemed transposable to humans^8^. Despite the recognition of its sexually-dimorphic activity, *in vivo* studies conducted so far on both male and female mice have focused on the impact of CAR activation on cytochromes P450 (CYPs) regulation^14^ or on genes involved in cell cycle and hepatocarcinogenesis^15, 16^. A genome-wide comparison of the effects of CAR activation in male and female mice is thus lacking.

Here, we used hepatic microarray and metabolomics in wild-type (*Car^+/+^)* and whole-body knock-out littermate (*Car^-/-^)* male and female mice treated either with Corn Oil (CO, vehicle) or TCPOBOP to elucidate the potential sex-dependent impact of CAR activation.

## Materials and Methods

### 1. Animals

*In vivo* studies were performed in a conventional laboratory animal room following the European Union guidelines for laboratory animal use and care. The current project was approved by an independent ethics committee (CEEA-86 Toxcométhique) under the authorization number 2019123014045837. The animals were treated humanely with due consideration to the alleviation of distress and discomfort. All mice were housed at 22°C ± 2°C on a 12 hours light (ZT0–ZT12) 12 hours dark (ZT12–ZT24) cycle and allowed free access to the diet (Teklad Global 18% Protein Rodent Diet) and tap water. ZT stands for Zeitgeber time; ZT0 is defined as the time when the lights are turned on. *Car^-/-^* mice (backcrossed on the C57BL/6J background) were engineered in Wei *et al.,* laboratory^1^ and are bred for 15 years in our animal facility. *Car^+/-^* mice were bred together and gave birth to true *Car^+/+^* and *Car^-/-^*littermate mice, which were then separated by sex and genotype at 4 weeks and randomly allocated to the different experimental groups. Nine-week-old male and female mice included in TCPOBOP groups received a daily intraperitoneal injection of 1,4-bis[2-(3,5- dichloropyridyloxy)]benzene (TCPOBOP, Sigma Aldrich) diluted in corn oil (CO, used as vehicle) at 3 mg/kg for 4 days while CO mice received corn oil only (Sigma Aldrich). One cage of n=6 mice per group was used. At ZT16 (6 hours after the last TCPOBOP injection), mice were anesthetized with isoflurane and xylazine (2%, 2 mg/kg) then blood from *vena cava* was collected into lithium heparin-coated tubes (BD Microtainer, Franklin Lake, NJ, USA). Plasma was prepared by centrifugation (1500 g, 15 min, 4°C) and stored at −80°C. Following euthanasia by cervical dislocation, liver and perigonadal white adipose tissue were removed, weighted and snap-frozen in liquid nitrogen and stored at −80°C.

To confirm *Fmo3* regulation upon TCPOBOP treatment another independent mouse experiment was conducted using the same experimental groups but a different TCPOBOP treatment timing leading to the same total dose of TCPOBOP: intraperitoneal injection either with TCPOBOP diluted in CO at 3 mg/kg every 2 days for 10 days or with CO. Mice were euthanized by cervical dislocation and liver was removed, weighted and snap-frozen in liquid nitrogen and stored at −80°C.

For thrombus formation analysis, another set of 7-week-old C57BL/6J females were purchased from Charles River laboratories, acclimatized for two weeks, then randomly allocated to the different experimental groups: Female Corn Oil (F CO, n=10), Female TCPOBOP (F TCPOBOP, n=10), (two cages per group). Then, mice were fed with 1% choline-enriched diet (D13090101, Research Diets) for 10 days. Mice included in TCPOBOP groups received an intraperitoneal injection of TCPOBOP diluted in corn oil at 3 mg/kg for the last 4 days of diet while CO mice received corn oil only. Whole blood was drawn from the inferior *vena cava* of anesthetized mice (100 mg/kg ketamine, 10 mg/kg xylazine) into heparin sodium (10 IU/ml) and mice were euthanized by cervical dislocation and liver was removed, weighted and snap-frozen in liquid nitrogen and stored at −80°C.

### 2. Gene Expression

Gene expression profiles were performed at the GeT-TRiX facility (GénoToul, Génopole Toulouse Midi-Pyrénées) using Agilent Sureprint G3 Mouse GE v2 microarrays (8x60K, design 074809) following the manufacturer’s instructions. Data acquisition and statistical analysis were performed as previously described^17^. A correction for multiple testing was applied using Benjamini-Hochberg procedure to control the False Discovery Rate (FDR). Probes with fold change (FC) ≥ 1.5 and FDR ≤ 0.05 were considered to be differentially expressed between conditions. The enrichment of Gene Ontology (GO) Biological Processes was evaluated using Metascape^18^. Data are available in NCBI’s Gene Expression Omnibus and are accessible through GEO Series accession number GSE228554.

For real-time quantitative polymerase chain reaction (RT-qPCR), 2 µg RNA samples were reverse-transcribed using the High-Capacity cDNA Reverse Transcription Kit (Applied Biosystems, Foster City, CA, USA). Additional file 1 presents the SYBR Green assay primers. Amplifications were performed using an ABI Prism 7300 Real-Time PCR System (Applied Biosystems, Foster City, CA, USA). RT-qPCR data were normalized to TATA-box-binding protein (*Tbp*) mRNA levels.

### 3. Proton Nuclear Magnetic Resonance (^1^H-NMR) Based Metabolomics

Plasma samples and liver polar extracts were prepared and analyzed using ^1^H- NMR-based metabolomics. All spectra were obtained on a Bruker DRX-600-Avance NMR spectrometer (Bruker) on the AXIOM metabolomics platform (MetaToul). Details on experimental procedures, data pre-treatment and statistical analysis were described previously^17^. Parameters of the final discriminating orthogonal projection on latent structure-discriminant analysis (O-PLS-DA) are indicated in the figure legends. To identify metabolites responsible for discrimination between the groups, the O-PLS-DA correlation coefficients (r^2^) were calculated for each variable. Correlation coefficients above the threshold defined by Pearson’s critical correlation coefficient (p< 0:05; |r|> 0.7; for n= 6 per group) were considered significant. For illustration purposes, the area under the curve of several signals of interest was integrated and significance tested with 2-way ANOVA as described below. For metabolite identification ^1^H-^13^C heteronuclear single quantum coherence (HSQC) spectra was obtained on one representative sample for each biological matrix. Lists of metabolites measured are presented in additional files 2 and 3.

### 4. Multi-omics analyses

Bidirectional correlations between plasma metabolites and hepatic transcripts were investigated using N-integration discriminant analysis with DIABLO, an algorithm that aims to identify a highly correlated multi-omics signature discriminating several experimental groups using the R package Mixomics v6.10.9^19^. We used 2 components in the models, and for the estimation of model parameters, the cross-validation procedure (CV) method was used. For the correlation networks, only correlations with a Spearman’s rank correlation coefficient > 0.96 were plotted.

### 5. Plasma Analyses

Alanine aminotransferase (ALT), phosphatase alkaline (ALP), total cholesterol, low-density lipoprotein (LDL-Cholesterol) and high-density lipoprotein (HDL-Cholesterol) were determined using a Pentra 400 biochemical analyzer (Anexplo facility, Toulouse, France).

### 6. Trimethylamine-N-oxide targeted LC-MS/MS measurement

For TMAO extraction method is described in details in additional methods.

### 7. Publicly available datasets and databases

Four independent gene expression datasets were found on the Gene Expression Omnibus data repository accessed in September 2019. GSE149229 compared hepatic transcriptome of humanized CAR mice (hCAR) fed a control diet or a phenobarbital (PB)-enriched diet. GSE98666 compared hepatic transcriptome of hCAR mice treated with CO, TCPOBOP or 6-(4-Chlorophenyl)imidazo[2,1-b][1,3]thiazole-5-carbaldehyde- O-(3,4-dichlorobenzyl)oxime (CITCO). GSE149228 and GSE57056 compared hepatic transcriptome of chimeric mice with a majority of human hepatocytes fed a control diet or a PB-enriched diet. Values for *Fmo3* gene expression were calculated using the GEO2R tool for microarray data and using GREIN^20^ for RNA sequencing data.

A list of drugs inducing sex-biased drug-induced liver injury was derived from the study conducted by George *et al.*^21^. For each drug, interaction with CYP2B6 was searched using DrugBank^22^ accessed in February 2023.

### 8. Thrombi formation under flow

Biochips microcapillaries (Vena8Fluoro+, Cellix) were coated with a collagen fibril suspension (50 µg/mL) and saturated with a solution of 0.5% bovine serum albumin in phosphate-buffered saline without Ca^2+^/Mg^2+^. Mouse blood was transferred into heparin (10 IU/mL), and DIOC6 (2 µM) was used to label platelets in whole blood. Using a syringe pump (PHD-2000; Harvard Apparatus) to apply a negative pressure, labeled blood was then perfused through a microcapillary for indicated time at a wall shear rate of 1500 seconds^-1^ (67.5 dynes/cm^2^). Thrombus formation was visualized with a X40 oil immersion objective for both fluorescent and transmitted light microscopy, light source was provided by Colibri (Zeiss) and was recorded in real time (1 frame every 20 seconds). Thrombi volumes are calculated by thresholding of surface covered by thrombi on slice of Z-stack images using IMARIS software^23^.

### 9. Other statistical analyses

All univariate statistical analyses were performed using GraphPad Prism version 9 (GraphPad Software, San Diego, CA). Outliers were identified using the ROUT method. The Kolmogorov–Smirnov test of normality was applied to all data. Two-way ANOVA was performed within each genotype using sex (male or female) and treatment (CO or TCPOBOP) as fitting factors for the models and P-value representing interactions are reported. If *P_sex*treatment_* was significant, Sidak’s multiple comparisons test was used as post-hoc test to determine which group differed from its appropriate control, otherwise P-values representing the main effects from the ANOVA model (namely sex or treatment) were reported. For platelet aggregation measures, a mixed-effect model was fitted using time and treatment as fixed effects. P<0.05 was considered as significant. Results are given as the mean ± SEM.

## Results

### Analysis of hepatic transcripts revealed a majority of sex-independent CAR-target genes upon TCPOBOP treatment

To investigate the potential sex-dependent consequences of CAR activation, we treated *Car^+/+^* and *Car^-/-^* male and female mice with TCPOBOP (Fig.1a). TCPOBOP did not affect total body, decreased perigonadal white adipose tissue and increased liver weights in *Car^+/+^* male and female mice (Fig. 1b). We confirmed *Car* deletion and observed increased expression of *Cyp2b10* and *Cyp2c55*, two prototypical CAR target genes, in TCPOBOP-treated *Car^+/+^* males and females. *Cyp2c55* induction by TCPOBOP was significantly higher in females (Fig. 1c). We characterized the impact of CAR activation by TCPOBOP on hepatic gene expression using microarrays. TCPOBOP was a very specific CAR agonist since there was no significantly regulated gene in *Car^-/-^* male mice, and only 2 in *Car^-/-^* female mice (Additional file 4), we thus continued our analysis using *Car^+/+^* mice only. Principal component analysis (PCA) of the entire expression data set from *Car^+/+^*mice revealed that individuals clustered separately according to treatment on the first axis and to sex on the second axis (Fig. 1d), illustrating a major effect of CAR activation on the liver transcripts. Comparison of the lists of differentially-expressed genes (DEG) upon TCPOBOP treatment in males *vs.* females *Car^+/+^*demonstrated that more than half of TCPOBOP-modulated genes were common between males and females (Fig. 1e, Additional file 5). Using all 4663 TCPOBOP-sensitive genes, we highlighted 2 gene clusters that exhibited sex-independent responses (Fig. 1f). Genes up-regulated upon TCPOBOP (cluster 1, 2366 genes) were mainly involved in “cell cycle” (*p=10^-^*^46^) and “cellular response to xenobiotic stimulus” (*p=10^-^*^25^, Fig. 1f and Additional file 6b), while down-regulated genes (cluster 2, 2297 genes) were enriched for “carboxylic acid catabolic process” (*p= 10^-^*^17^) and “regulation of lipid metabolic process” (*p=10^-^*^14^, Fig. 1f and Additional file 6d). We focused on well-described CAR target genes involved in cell cycle (Fig. 1g) and carbohydrate and lipid metabolism (Fig. 1h) and confirmed that regulation upon TCPOBOP treatment was similar in both males and females (Additional file 7).

**Figure 1:**
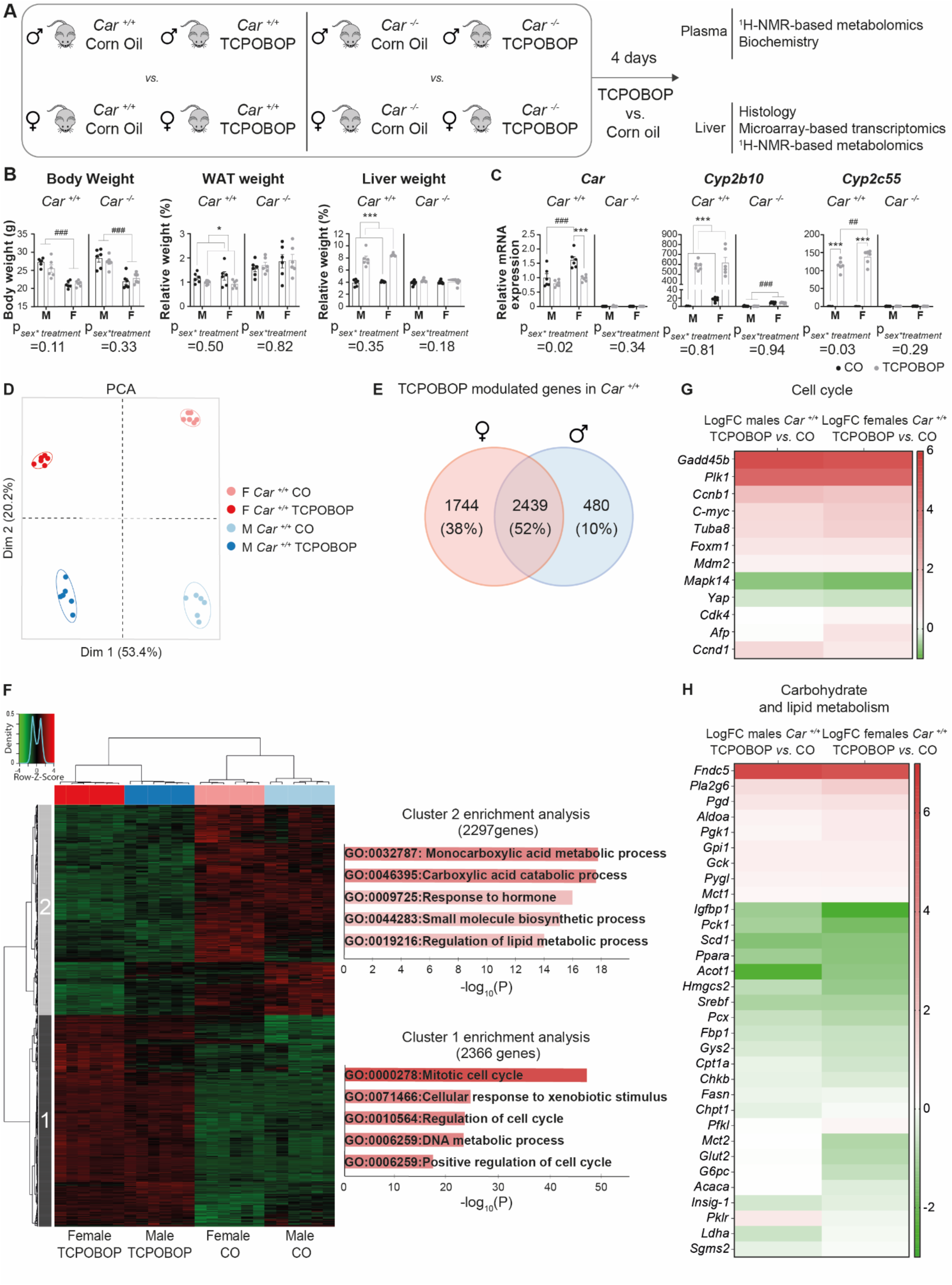
Modulation of classical CAR-controlled pathways is sex-independent. *A) Experimental design. B) Body weight, perigonadal WAT and liver weights. C) Hepatic gene expression. D) PCA of the whole liver transcriptomic dataset in Car^+/+^ mice. E) Venn diagram representing the number of genes significantly modulated by TCPOBOP in the liver of Car^+/+^ male and female mice. F) Hierarchical clustering and pathway enrichment analysis using the 4663 genes significantly regulated upon TCPOBOP in males or females. G-H) Heatmaps representing the log(fold-change) of gene expression between TCPOBOP- and CO-treated Car^+/+^ males and females for genes involved in cell cycle. (G) and carbohydrate and lipid metabolism. (H) Results are given as the mean ± SEM *treatment effect, #sex effect. * or # p<0.05, ** or ## p<0.01, *** or ### p<0.001.*

### Role of CAR in sexually dimorphic regulation of hepatic gene expression in response to TCPOBOP

TCPOBOP impacted a much higher number of genes in the liver of females *vs.* males (about 40% more DEG in females *vs.* males, Fig. 1e). Accordingly, PCA of microarray data projected on the 2^nd^ and 3^rd^ principal components showed a distinct clustering of TCPOBOP- and CO-treated females while males from both groups were merged (Fig. 2a). To identify genes with sex-dependent regulation upon TCPOBOP, we next focused on the DEG with a significant interaction between sex and treatment (Additional file 8). These 486 sex-specific DEG clustered within 4 distinct expression profiles (Fig. 2b and 2c). Female-specific up-regulated genes (cluster 1, 215 genes) were involved in xenobiotic metabolism (*p=10^-^*^12^) and extra-cellular matrix remodeling (*p=10^-^*^6^) and contained genes encoding for collagens (*Col4a1*), extra-cellular matrix degrading metalloproteinases (*Mmp12, Mmp13*) and proteinases involved in the processing of procollagens (*Adamst2, Adamst4, Adamst14* and *Adamst15*), while female-specific down-regulated genes (cluster 3, 170 genes) were involved in phase I xenobiotic metabolism (*p=10^-^*^8^) and FMO-dependent oxidations (*p=10^-^*^6^), while female-specific (Fig. 2d-f). As previously seen, male specific CAR target genes were fewer. The 37 male-specific up-regulated genes (cluster 2, 37 genes) were involved in “steroid metabolism” (*p=10^-^*^5^*^.^*^5^), while no significantly enriched metabolic pathway was found using the 64 male-specific down-regulated genes (cluster 4, 64 genes) (Fig. 2d-e and Additional file 8). Overall, this analysis provided evidence that 10% of the TCPOBOP- sensitive genes were regulated in a sex-dependent way (Fig. 2g).

**Figure 2:**
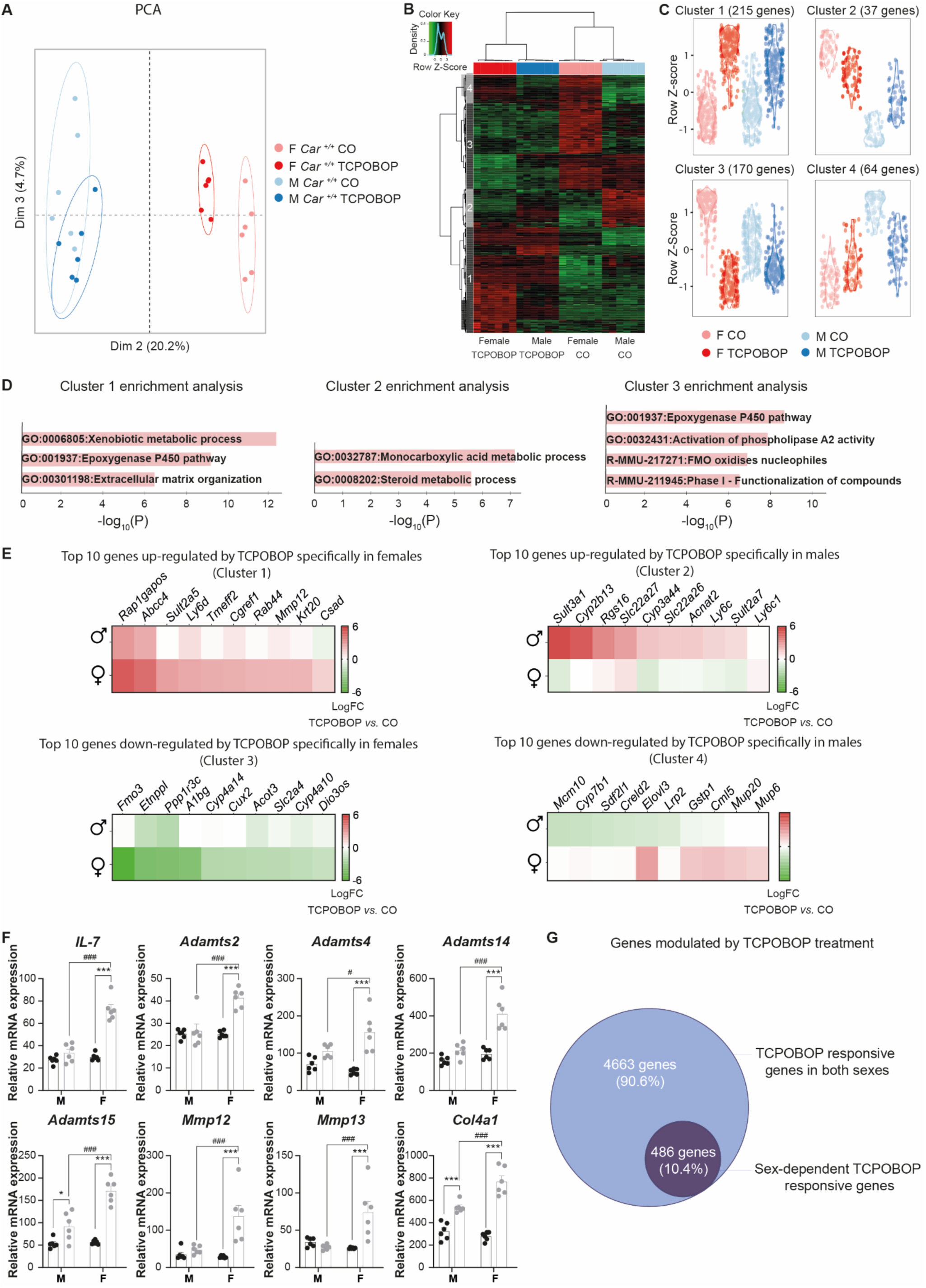
Identification of sex-dependent CAR-sensitive genes. *A) PCA of the liver transcriptomic dataset in Car^+/+^ mice. B) Hierarchical clustering using the 486 genes with significant P_sex*treatment._ C) Gene expression profiles in each cluster. D) Pathway enrichment analysis. E) Heatmaps representing the log(fold-change) of gene expression between TCPOBOP- and CO-treated Car^+/+^ males and females for the top 10 genes in each cluster. F) Hepatic gene expression of genes involved in inflammation and fibrosis derived from microarray data. G) Genes modulated by TCPOBOP treatment in a sex-dependent and –independent way.*

### CAR regulates plasma lipoprotein metabolism in a sex-independent manner

We next explored the systemic consequences of CAR activation using plasma metabolomics. PCA analysis of the whole plasma metabolic profiles showed a separation of male *vs.* female mice on the first principal component, illustrating a constitutive sexual dimorphism in plasma metabolite levels, while TCPOBOP-treated *Car^+/+^* were discriminated from *Car^-/-^*mice and from *Car^+/+^* CO-treated mice on the second principal component, illustrating a significant effect of TCPOBOP on plasma metabolites (Fig. 3a). As seen for hepatic transcripts, there was no significant difference between metabolic profiles from TCPOBOP *vs.* CO-treated *Car^-/-^* mice (Additional file 9). In *Car^+/+^* males, TCPOBOP treatment decreased glucose, increased lactate levels and had a major impact on circulating lipoproteins: cholesterol and several broad lipid peaks were strongly decreased upon TCPOBOP treatment, while other lipid peaks were increased (Fig. 3b). In *Car^+/+^* females, cholesterol and lipid signals were changed in a similar manner than in males (Fig. 3c). Putative assignment of these differential peaks revealed that the decreased cholesterol peak mainly reflected HDL-cholesterol, decreased lipid peaks mainly belonged to LDL and increased lipids belonged to very low-density lipoproteins (VLDL). Area under the curve for selected HDL-cholesterol, lipid-LDL and lipid-VLDL signals further illustrated this CAR-dependent impact of TCPOBOP treatment on circulating lipoproteins in both sexes (Fig. 3d). The observed decrease in total plasma-, LDL- and HDL-cholesterol were confirmed through additional biochemical assays (Additional file 10).

**Figure 3:**
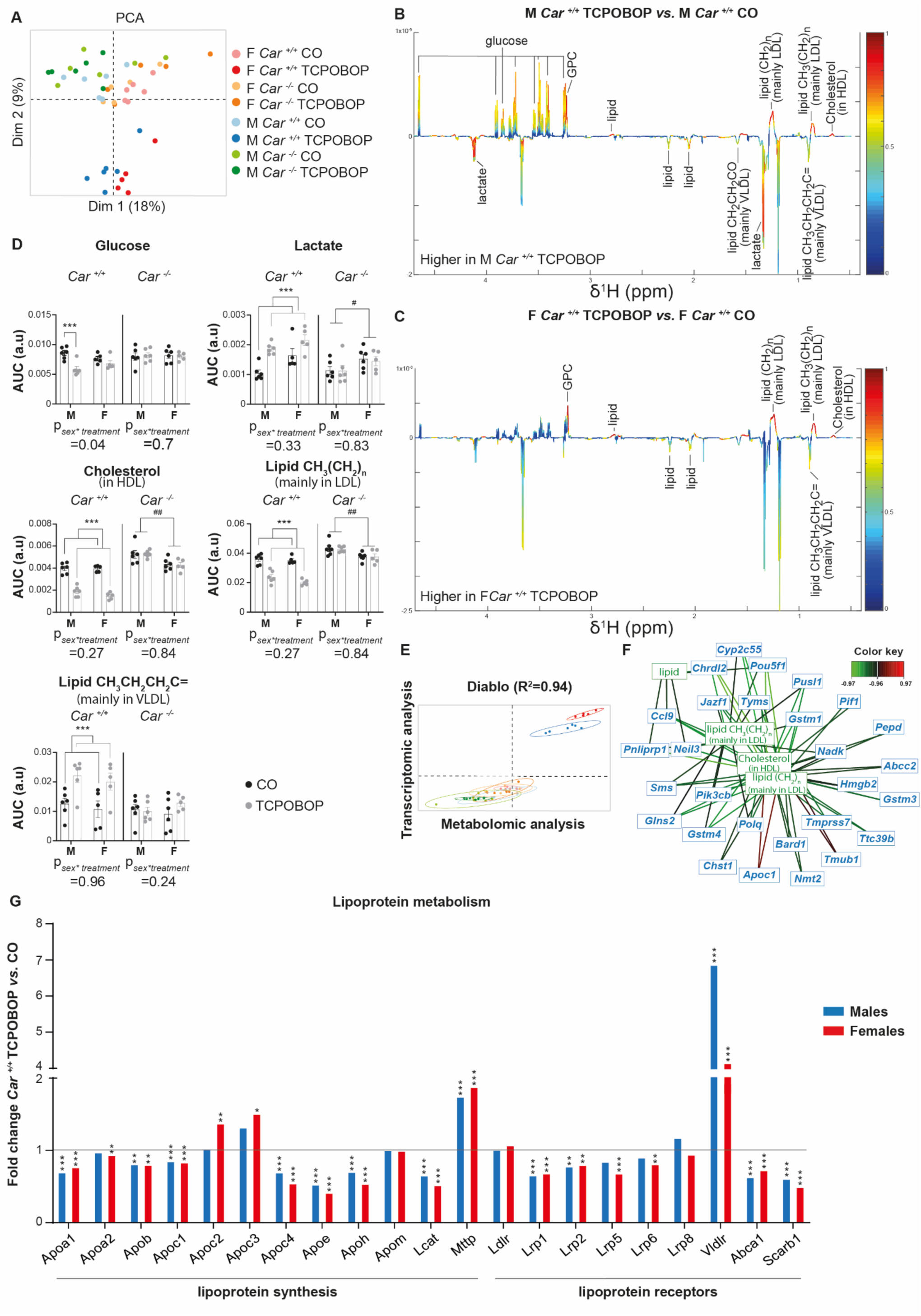
Sex-independent impact of CAR activation on lipoprotein metabolism. *A) PCA of plasma metabolomic dataset. B) Coefficient plots related to the PLS-DA models discriminating between plasma spectra from TCPOBOP vs. CO Car^+/+^ males. Parameters of the PLS-DA model: Q^2^Y=0.84, p=0.001. C) Coefficient plots related to the PLS-DA models discriminating between plasma spectra from TCPOBOP vs. CO Car ^+/+^ females. Parameters of the PLS-DA model: Q^2^Y=0.89, p=0.001. D) AUC of the ^1^H-NMR spectra was integrated for glucose, lactate, cholesterol, lipid CH3(CH2)n (mainly in LDL), lipid CH3CH2CH2C= (mainly in VLDL) signals. E) Multi-omic integrative analysis performed on plasma metabolomic and hepatic transcriptomic datasets. F) Correlation network between hepatic transcripts and plasma metabolites. G) Fold- change (TCPOBOP- vs. CO-treated Car^+/+^ mice) of hepatic expression for genes involved in lipoprotein metabolism. *treatment effect, #sex effect. * or # p<0.05, ** or ## p<0.01, *** or ### p<0.001.*

Plasma metabolomics and hepatic transcriptomic data statistical integration revealed a strong correlation between plasma metabolites and liver transcripts (R^2^=0.94) regardless of sex (Fig. 3e). Correlation network highlighted a strong positive correlation between *Apoc1* mRNA and plasma LDL-cholesterol and HDL-cholesterol (Fig. 3f). This led us to investigate more closely the hepatic expression of genes involved in hepatic cholesterol metabolism (Fig. 3g). CAR activation significantly decreased the expression of the majority of apolipoproteins. Expression of the *Ldl receptor* (*Ldlr*), which is responsible for LDL clearance was unchanged, however, that of *Vld receptor* (*Vldlr*) was increased by a factor of 4 upon TCPOBOP. *Mttp*, the protein that transports triglycerides and cholesterol esters in the endoplasmic reticulum for VLDL synthesis was also significantly increased. Moreover, TCPOBOP treatment impacted cholesterol and bile acid metabolism with decreased expression of genes involved in cholesterol transport, decreased expression of genes involved in bile acid synthesis and increased expression of genes involved in bile acid detoxication and transport. All significant changes in hepatic mRNA and plasma metabolites related to cholesterol metabolism were CAR-dependent and were similar in both sexes (Additional file 11 and 12). Altogether, these results illustrate that CAR activation deeply modulates hepatic and systemic cholesterol metabolism in a sex-independent way.

### TCPOBOP induces liver oxidative stress and toxicity in a sex-biased way

We next performed metabolic profiling of hydrophilic metabolites in liver tissue. PCA of entire metabolic profiles revealed a distinct clustering of *Car^+/+^* mice treated with TCPOBOP *vs.* all other mouse groups on the first principal component, while male and female mice were separated on the 2^nd^ component, revealing once again a major impact of CAR activation on liver metabolites (Fig. 4a). TCPOBOP-treated males displayed significant changes in hepatic levels of many amino-acids (increased glutamine and glutamate and decreased valine, leucine and isoleucine), energy-related metabolites (increased lactate and 3-hydroxybutyrate and decreased succinate), cell membrane constituents (decreased choline and glycerophosphocholine and increased phosphocholine) and metabolites involved in oxidative-stress (decreased hypotaurine) (Fig. 4b). In females, most of these metabolites followed the same pattern, with perturbations of metabolites involved in oxidative stress being more pronounced than in males (significant increased levels of reduced (GSH), oxidized (GSSG) and total (Gsx) glutathione) (Fig. 4c). AUC for glutathione signals illustrated that TCPOBOP induced a more pronounced hepatic oxidative stress in females than in males (Fig. 4d). All significant changes in hepatic metabolites related to oxidative stress upon TCPOBOP were CAR-dependent (Additional file 13). Finally, sex-dependent TCPOBOP hepatic toxicity was confirmed through biochemical quantification of plasmatic markers (Fig. 4e-f). Circulating levels of alanine aminotransferase (ALT) were significantly higher in TCPOBOP-treated *Car^+/+^* females *vs*. males, while plasma alkaline phosphatase (ALP) was significantly increased in both sexes upon TCPOBOP with a tendency to higher levels in males (*P_sex*treatment_=0.12*).

**Figure 4:**
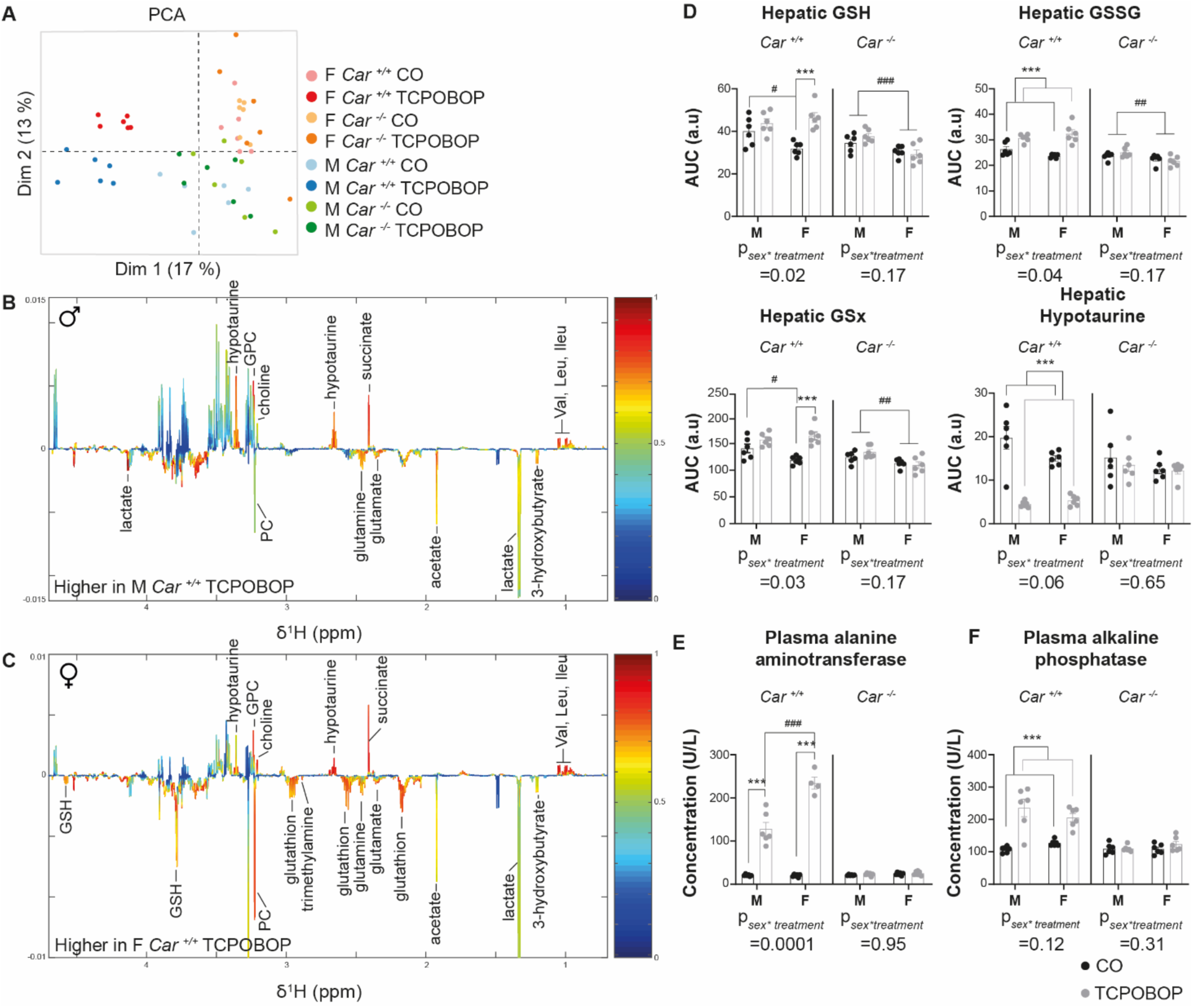
TCPOBOP treatment increases liver oxidative stress and toxicity in a sex-biased way. *A) PCA of the liver metabolomic dataset. B) Coefficient plots related to the PLS-DA models discriminating liver extract spectra from TCPOBOP- and CO-treated Car ^+/+^ males. Parameters of the PLS-DA model: Q^2^Y=0.89, p=0.001. C) Coefficient plots related to the O-PLS-DA models discriminating between liver extract spectra from TCPOBOP- and CO-treated Car ^+/+^ females. Parameters of the PLS-DA model: Q^2^Y=0.91, p=0.001. D) AUC of the ^1^H-NMR spectra was integrated for the glutathione signals (GSH, reduced form; GSSG, oxidized form; GSx, total glutathione) and for hypotaurine. E) Plasma alanine aminotransferase. F) Plasma alkaline phosphatase. Results are given as the mean ± SEM. *treatment effect, #sex effect. * or # p<0.05, ** or ## p<0.01, *** or ### p<0.001.*

We further investigated the potential role of CAR in sex-biased hepatic toxicity in humans using a previously published list of drugs inducing drug-induced liver injury (DILI)^21^. George *et al.*^21^ conducted a data-mining analysis of the WHO VigiBase^TM^ and calculated the reporting frequency of liver events for 375 drugs with DILI potential. They reported a list of 17 drugs with man-biased reporting frequency and a list of 41 drugs with woman-biased reporting frequency (Table 1). We wondered whether drugs from these lists were potentially interacting with CAR and used interaction with CYP2B6 as a proxy for CAR activity as it was described as CAR primary target in humans^24^. Among the 17 drugs with man-biased reporting frequency, we found only 3 of them were described as interacting with CYP2B6 (3/17=17,6%), while this proportion reached 16/40 (40%) in the woman-biased list of drugs (*p=0.0507* using the Chi-square test). Overall, these results demonstrate a sex-dependent impact of CAR activation on liver metabolism and toxicity profile, which might be relevant also in human DILI.

**Table 1.** Drugs triggering sex-dependent Drug Induced Liver Injury (DILI) and their pharmacological interaction with CYP2B6.

| Drugs with male-biased reporting frequency | Interaction with CYP2B6 | Drugs with female-biased reporting frequency | Interaction with CYP2B6 |
| --- | --- | --- | --- |
| Alclofenac |  | Amitriptyline | substrate |
| Amoxicillin |  | Amlodipine | substrate |
| Azithromycin |  | Atorvastatin | inducer |
| Clarithromycin |  | Azathioprine |  |
| Clindamycin |  | Bosentan |  |
| Cyclophosphamide | inducer | Clomipramine |  |
| Cyproterone |  | Clopidogrel | substrate |
| Danazol |  | Clozapine |  |
| Doxorubicin | inhibitor | Diclofenac | substrate |
| Estradiol |  | Disulfiram |  |
| Etanercept |  | Duloxetine | inhibitor |
| Isotretinoin |  | Efavirenz | substrate inducer |
| Methyltestosterone | substrate | Erlotinib |  |
| Stanozolol |  | Erythromycin |  |
| Succimer |  | Ethambutol |  |
| Trimethoprim |  | Flucloxacillin |  |
| Triptorelin |  | Haloperidol |  |
| **TOTAL** | **3/17=17.6%** | Halothane | substrate |
|  |  | Ibuprofen |  |
|  |  | Imatinib |  |
|  |  | Interferon Beta-1a |  |
|  |  | Isoniazid |  |
|  |  | Ketoconazole | inhibitor |
|  |  | Leflunomide |  |
|  |  | Lisinopril |  |
|  |  | Metoprolol |  |
|  |  | Nitrofurantoin |  |
|  |  | Olanzapine |  |
|  |  | Paclitaxel |  |
|  |  | Paroxetine | inhibitor |
|  |  | Phenytoin | inducer |
|  |  | Pyrazinamide |  |
|  |  | Ramipril |  |
|  |  | Risperidone |  |
|  |  | Ritonavir | inducer |
|  |  | Sevoflurane | substrate |
|  |  | Sulfasalazine |  |
|  |  | Ticlopidine | inhibitor |
|  |  | Troglitazone | inducer |
|  |  | Valproic Acid | sustrate |
|  |  | **TOTAL** | **16/40=40%** |

### CAR modulates liver trimethylamine metabolism through regulation of *Fmo3* gene expression mostly in females

Another intriguing sex-dependent hepatic impact of TCPOBOP was the female-specific, CAR-dependent increased level of trimethylamine (TMA) (Fig. 4c, Fig. 5a) TMA is a gut-microbiota dependent metabolite that is metabolized to trimethylamine N-oxide (TMAO) by the liver-specific flavin monooxygenase 3 (FMO3)^25^. This result was in accordance with the female-specific decrease of *Fmo3* mRNA expression observed previously in the microarray data (Fig. 2e) and confirmed here with RT-qPCR (Fig. 5b). Finally, we quantified hepatic TMAO and confirmed a 2 fold-decrease of this metabolite in TCPOBOP-treated *Car^+/+^* female compared to vehicle-treated females (Fig. 5c). Female-biased inhibition of *Fmo3* mRNA by CAR activation was reproducible in an independent study in which mice were treated with TCPOBOP or CO every two days for 10 days (Fig. 5d). Next, we analyzed *Fmo3* hepatic expression in several publicly available gene expression datasets. In the first experiment, *Car^-/-^* mice were knocked-in with human CAR coding sequence^26^ and were fed diets containing 0 (CTRL) or 1000ppm phenobarbital (PB), an indirect activator of both mCAR and hCAR^27^. We found that PB-fed hCAR mice had significantly lower expression of hepatic *Fmo3* mRNA compared to control mice (Fig. 5e). The second study compared wild-type (WT) mice treated with TCPOBOP *vs.* CO-treated mice and hCAR mice treated with CITCO (a specific agonist of hCAR) *vs.* CO-treated hCAR mice^28^. Both TCPOBOP and CITCO-treated mice had significantly lower *Fmo3* hepatic mRNA compared to their relative controls (Fig. 5f). Finally, the last two studies were conducted in chimeric mice with human hepatocytes treated with PB^27, 29^. Again, we found a significant decrease in *Fmo3* hepatic gene expression in response to PB in both datasets (Fig. 5g and 5h). It is worth noting that all publicly available studies were conducted in male mice only, which could explain why the decrease in *Fmo3* expression seen upon CAR activation by PB or CITCO was of lower magnitude than that seen in our own *in vivo* experiments in females. Finally, we took advantage of the only available ChIP-seq analysis of hCAR binding *in vivo* to date^30^ and observed that, among the 6364 unique genes associated with high-confidence hCAR-binding genes, *Fmo3* was found as a hCAR-direct binding gene with a *p-value=1,26 10^-^*^29^ (Fig. 5j). Thus, in females, CAR activation by TCPOBOP perturbated the metabolism of TMAO from TMA by inhibiting the expression of *Fmo3* (Fig. 5i) and this result might be relevant to humans.

**Figure 5:**
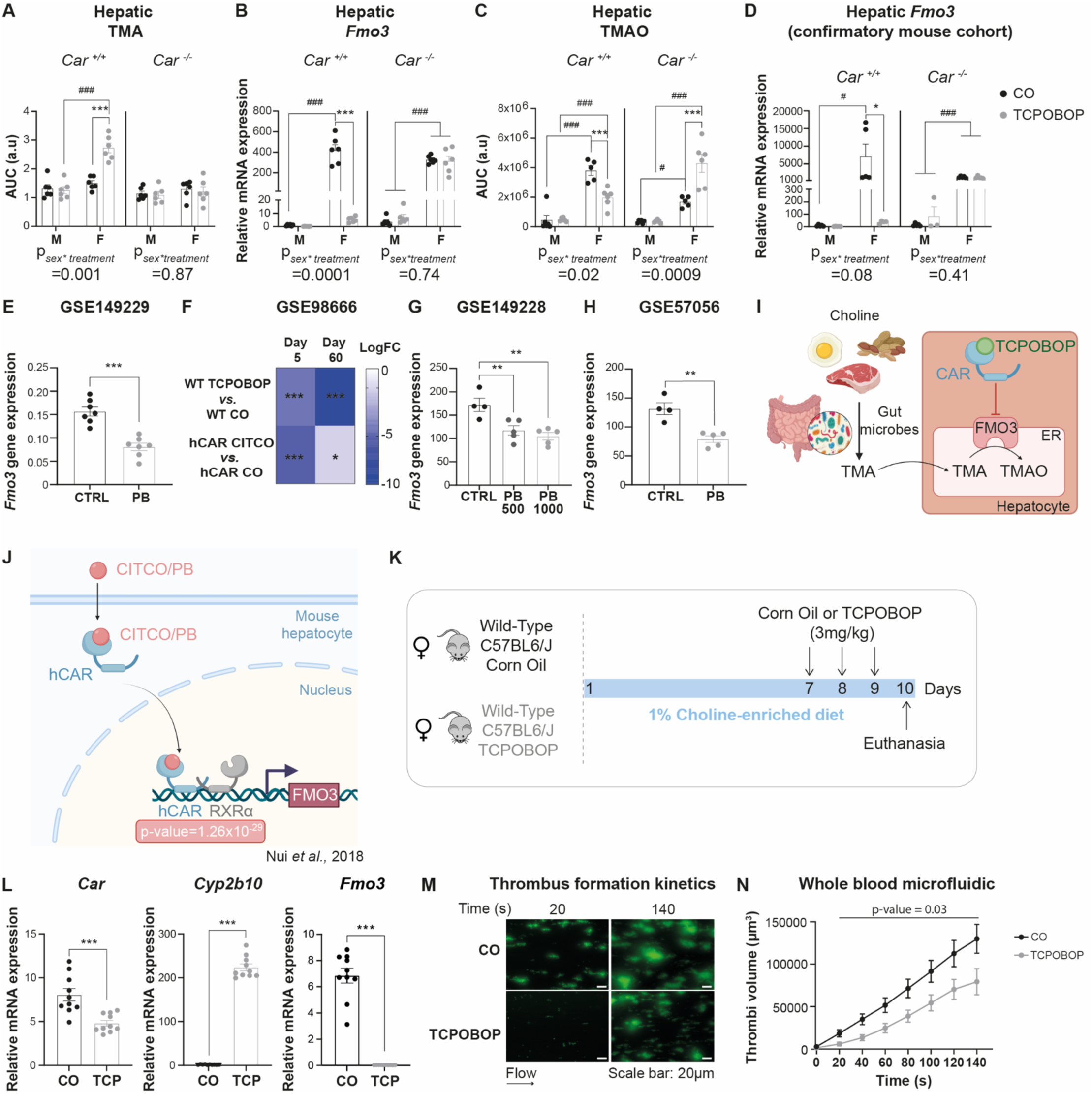
CAR modulates liver TMA metabolism by regulating Fmo3 gene expression, mostly in females. **(A)** *AUC of the ^1^H-NMR spectra for TMA. B) Hepatic gene expression. C) Hepatic content of TMAO. D) Hepatic gene expression in independent confirmatory experiment. E-H) Hepatic mRNA expression derived from publicly available datasets*. • *Proposed impact of CAR activation on TMA metabolism (Created with BioRender.com). J) Proposed model for direct binding of hCAR to Fmo3 regulatory DNA sequences based on* ^30^ *(Created with BioRender.com). K) Experimental design*. *L) Hepatic gene expression. M) Representative images of platelet adhesion. N) Quantification of platelet adhesion to a microfluidic chip surface. Results shown are mean ± SEM. *treatment effect, #sex effect. * or # p<0.05, ** or ## p<0.01, *** or ### p<0.001*.

### CAR activation decreases platelet hyperactivity induced by dietary choline supplementation

FMO3 activity and TMAO levels have been shown to modulate platelet hyperresponsiveness and thrombosis potential^31, 32^. We thus wondered whether CAR activation could also impact platelet function. To enhance platelet responsiveness, female mice were fed a choline-enriched diet before treatment with TCPOBOP (Fig. 5k). We confirmed significant hepatic CAR activation (Fig. 5l) and examined thrombi formation. Results showed that whole blood from mice treated with TCPOBOP formed smaller thrombi over time compared to blood from control mice. Thus, *in vivo* TCPOBOP treatment modulates platelet activation and reduces the thrombotic risk of these mice (Fig. 5m and 5n).

## Discussion

The liver appears to be one of the most sexually-dimorphic organ and expression of genes involved in drug metabolism is sexually dimorphic in rodents and humans^8, 33^. CAR is the target of many drugs and is widely involved in the control of expression of xenobiotic metabolizing genes. However, a genome-wide description of sex-specific CAR-dependent sensitive genes was lacking. Here, we provide an exhaustive study of the transcriptomic impact of acute pharmacological CAR activation in male *vs.* female mice and novel insights into the metabolic impact of this activation.

First, the majority of TCPOBOP-modulated genes was regulated in a similar manner in male and female livers, especially those involved in cell cycle. Our results are consistent with other studies revealing that chronic activation of CAR using TCPOBOP promotes tumor formation in rodents in a CAR-dependent way^34–36^. The underlying mechanisms depend on direct CAR-dependent induction of *Mdm2*, a primary inhibitor of P53 and induction of the transcription factor *FoxM1*^37^, which is essential for initiation of carcinogen-induced liver tumors, thus resulting in modulation of many genes implicated in cell proliferation, cellular growth, apoptosis and cell differentiation, such as *Gadd45b, Ccnd1, Ccnb1, C-myc, and Yap*^37, 38^. Few studies have highlighted that sex could influence TCPOBOP-induced liver proliferation but showed inconsistent results. Some studies described that female mice were more sensitive to TCPOBOP-induced liver proliferation^12, 39^ while others showed no tumor development in female mice treated with the genotoxin diethylnitrosamine (DEN) followed by TCPOBOP, compared to males^40^. Similarly, after a single injection of TCPOBOP, male mice displayed a deeper disturbance of key cell cycle genes^16^. Unlike these studies, we did not reveal any sexual dimorphism on hepatomegaly and cell cycle gene modulation upon CAR activation. However, a long-term analysis of TCPOBOP-induced liver tumors conducted in male and female mice in parallel would be required to further investigate this discrepancy.

We next observed a strong impact of TCPOBOP treatment on plasma lipoproteins, with decreased total-, HDL- and LDL-cholesterol measured in both sexes. This result is consistent with previous findings describing that TCPOBOP decreased circulating levels of plasma HDL in both WT and transgenic mice expressing human Apolipoprotein A-1, at least in part through down-regulation of *ApoA-1* hepatic gene expression^41^. Similarly, in *Ldlr^-/-^* mice fed a western-diet TCPOBOP decreased circulating levels of ApoB-containing lipoproteins (mainly VLDL and LDL) and reduced the development of atherosclerotic lesions^42^, presumably through CAR-mediated induction of the VLDL-receptor, a receptor involved in VLDL and LDL clearance as a backup for the LDL receptor^43^. Here, we confirm that CAR activation results in decreased *ApoA-1* and increased *Vldlr* hepatic mRNA levels, which could both participate to the observed decrease of plasmatic HDL and LDL levels. Moreover, we also observed a strong decrease in hepatic expression of other major lipoprotein-coding genes (namely *ApoB, Apoc1* and *ApoE*), and of the *Lecithine Cholesterol Acyl Transferase* (*Lcat*, another constitutive component of HDL) which could also play a role.

Another well-known function of CAR is its ability to promote bile acid detoxification during cholestasis^44, 45^. As previously described, we found that the expression of genes involved in hydroxylation, sulfation, and excretion of bile acids was significantly enhanced upon CAR activation, while expression of genes involved in bile acid synthesis and cholesterol transport was decreased in both sexes. The emerging role of CAR in cholesterol homeostasis represents new perspectives in the treatment of hypercholesterolemia and atherosclerosis^41, 46^. The current study confirms and extends these previous studies reporting the effects of TCPOBOP on hepatic expression of genes involved in bile acid, cholesterol and lipoprotein metabolism, as well as those on lipoprotein concentrations, are sex-independent, at least in mice.

Our findings may have clinical relevance. A recent study combining genome-wide analysis of cholestatic mice genetic-models and data-mining of human patient cohorts with various liver diseases unraveled a significant enrichment of CAR-sensitive genes in cholestatic livers specifically^47^. Moreover, authors demonstrated that CAR activation in cholestasis led to alterations of drug metabolism with significant effects on drug-induced hepatotoxicity. Drug-induced liver injury (DILI) is still a serious clinical concern and one of the most common drug adverse reaction. DILI clinical phenotype appears to be influenced by age and gender^48, 49^. Here, we found that, upon TCPOBOP, 385 genes displayed a female-specific *vs.* 101 genes with a male-specific response. Many female-specific genes were involved in extra-cellular matrix organization. We also observed stronger perturbations of hepatic metabolites involved in glutathione metabolism in female’s livers compared to males, reflecting higher hepatic oxidative stress. Finally, we highlighted a sexually-dimorphic impact of CAR activation on clinical markers of liver toxicity, namely significantly higher levels of alanine aminotransferase in females compared to males, and a trend toward higher alkaline phosphatase levels in males. This result is consistent with the sex-influence on DILI clinical phenotype with cytolytic damage being more frequently observed in women while cholestatic damage presented a man predominance^48, 49^. We also found a significant enrichment of drugs described as CYP2B6 substrates and/or inducers within the drugs with women-biased DILI reporting frequency compared to those with men-biased DILI reporting frequency. It is well known that women experience higher rates^50^, and more severe^51^ adverse drug reactions than men do. However, mechanistic explanations for these observations are often lacking. Our present results suggest that drugs interacting with CAR may be considered with particular attention before their use in women. However, limitations of our study include the use of only one drug (TCPOBOP) while DILI has been shown to depend both on patient characteristics and drug properties^21^. Our results therefore need to be confirmed with other drugs that act as CAR agonists.

Another novel finding from our study was the strong increase in hepatic TMA upon TCPOBOP observed in female mice specifically. TMA is a product of the gut microbial metabolism of phosphatidylcholine, choline and L-carnitine. It is transported from the gut to the liver *via* the portal vein and N-oxidized into TMAO by host FMO3^52^. Analysis of natural genetic variation in inbred strains of mice indicate that FMO3 and TMAO are significantly correlated and explain more than 10% of the variation in atherosclerosis^52^. Since then, it has been confirmed that high circulating levels of TMAO are linked to increased thrombotic and cardiovascular risks in animal and human studies, even after adjustment for known cardiovascular risk factors^53, 54^. Consistent with increased TMA, hepatic *Fmo3* mRNA expression and TMAO concentration were both strongly decreased in *Car^+/+^* females treated with TCPOBOP. We confirmed the CAR-dependent regulation of *Fmo3* mRNA in publicly available cohorts that used different hCAR models and different mCAR and hCAR agonists, therefore suggesting that the regulation of *Fmo3* expression is not dependent on the CAR agonist used and might be relevant in humans. In rodents, hepatic *Fmo3* knockdown was sufficient to decrease diet-dependent platelet responsiveness and thrombotic potential^31, 32^. Here, we observed that, in conditions of diet-induced platelet hyperresponsiveness, CAR activation was indeed sufficient to significantly modulate thrombus growth in female mice. We postulate that this effect is, at least in part, due to the CAR-mediated down-regulation of *Fmo3* expression and activity. Nowadays, drugs represent the main cause of platelet dysfunction^55, 56^. Our results suggest that drugs or other xenobiotics (pollutants, foods…) that interact with CAR decrease thrombus formation in a pro-thrombotic context. These compounds may provide a beneficial effect by modulating platelet activation and thrombosis. We therefore, highlight a new axis between hepatic xenobiotic metabolism and blood hemostasis. We suggest that this axis may be especially relevant in women. However, there are important species-specific sex-differences in *FMO3* expression: its expression is female-specific in mice due to modulation by sex steroids^57^, while its abundance was significantly associated with female in humans but to a much lower extend than in rodents^58^. Thus, the gender-specificity and clinical relevance of this CAR-FMO3- TMAO-platelet axis deserves further investigation. In addition to platelet function and thrombotic risk, an increase in the TMAO plasma concentration has also been shown to increase the risk of impaired glucose tolerance^59^, colorectal cancer^60^, chronic kidney disease^61^ and overall mortality^62^. Whether drugs interacting with CAR could also influence these TMAO-dependent endpoints deserves further investigations.

In summary, the present study provides an exhaustive description of the sex-independent and sex-dependent CAR-sensitive genes and demonstrates a stronger impact of CAR pharmacological activation on female’s hepatic transcriptome and metabolism. Additionally, CAR activation impacted the TMA-FMO3-TMAO pathway in females, which might link drugs and environmental xenobiotic exposure with platelet aggregation and other TMAO-sensitive physiological responses.

## Abbreviations

ALP,: alkaline phosphatase;
ALT,: alanine aminotransferase;
CAR,: constitutive androstane receptor;
ChIP-seq,: chromatin immunoprecipitation sequencing;
CITCO,: 6-(4-Chlorophenyl)imidazo[2,1-b][1,3]thiazole-5-carbaldehyde-O-(3,4-dichlorobenzyl)oxime;
CO,: corn oil;
CV,: cross-validation procedure;
CYPs,: cytochromes P450;
Cy3,: cyanine3;
DEG,: differentially expressed genes;
DEN,: diethylnitrosamine;
DILI,: drug-induced liver injury;
DIOC6,: 3,3’-dihexyloxacarbocyanine iodide;
FC,: fold change;
FDR,: false discovery rate;
Fmo3,: flavin monooxygenase 3;
GEO,: gene expression omnibus;
GO,: gene ontology;
GSH,: reduced glutathione;
GSSG,: oxidized glutathione;
GSx,: total glutathione;
hCAR,: humanized constitutive androstane receptor;
HSQC,: heteronuclear single quantum coherence;
Lcat,: lecithine cholesterol acyl transferase;
Ldlr,: low-density lipoprotein receptor;
mCAR,: murine constitutive androstane receptor;
MRM,: multiple reaction monitoring;
Mttp,: microsomal triglyceride transfer protein;
O-PLS-DA,: orthogonal projection on latent structure-discriminant analysis;
PB,: phenobarbital;
PCA,: principal component analysis;
RT-qPCR,: real-time quantitative polymerase chain reaction;
Tbp,: TATA-box-binding protein;
TCPOBOP,: 1,4-bis[2-(3,5-dichloropyridyloxy)] benzene;
TMA,: trimethylamine;
TMAO,: trimethylamine N-oxide;
Vldr,: very low-density protein receptor;
WAT,: perigonadal white adipose tissue;
WT,: wild-type;
ZT,: zeitgeber time;
^1^H-NMR,: proton nuclear magnetic resonance.

## Acknowledgements

We thank all members of the EZOP (Experimental Zootechny), UMR Toxalim, Toulouse, France, staff for their careful help with this project. We thank the staff from the Genotoul: Anexplo, Get-TriX and Metatoul-AXIOM facilities.

