## Additional files for "Pharmacological activation of constitutive androstane receptor induces female-specific modulation of hepatic metabolism"

**Additional files and methods for:**

Toxalim (Research Centre in Food Toxicology)

### Additional methods

1. Trimethylamine-N-oxide targeted LC-MS/MS measurement

For TMAO extraction method from [1] was modified. Liver tissue was combined with 425 µl ice-cold methanol and 75 µl 1% formic acid. Samples were homogenized using a TissueLyser II (30 Hz, 10 min; Retsch Qiagen). After centrifugation (5 min, 8°C, 14.000 rpm) 170 µl of supernatant was transferred to a new tube and evaporated to dryness (Eppendorf concentrator plus, Eppendorf). Prior to measurement, samples were resuspended in LC solvent A:B (1:1), volume depending on the liver weight used for extraction. Of each sample, 10 µl were injected onto an Agilent 1290 II infinity UPLC system (Agilent Technologies Inc., Santa Clara, USA) coupled online to a QTRAP®6500 mass spectrometer (Sciex, Framingham, USA). Chromatographic run was achieved using a reverse phase XSelect HSS T3 XP column (2.1 x 150 mm, 2.5 µm, 100 Å; Waters, Milford, USA) with a binary solvent system 5% acetonitrile with 0.1% formic acid in water (A) and 5% water with 0.1% formic acid in acetonitrile (B). Constant flow rate was set to 250 µl and column oven temperature was at 30°C. Mass spectrometric measurement was performed in positive ionization mode. For identification and quantitation, a scheduled multiple reaction monitoring (MRM) method was used with m/z 76.0 → 59.0 and m/z 75.0 → 42.2 as specific transitions for TMAO. Peak areas were determined in Analyst® Software (v. 1.7.1, Sciex) and exported. Analytical standard trimethylamine N-oxide (TMAO) was purchased from Sigma Aldrich (St Louis, MO, USA). All solvents for MS were of analytical grade purity. Experimental water (resistivity of 18.2 MΩ cm) was purified using a Milli-Q system (Millipore, Milford, MA, USA).

### Additional files

**Additional File 1:** Primers.

| Gene | NCBI Refseq | Forward primer | Reverse primer |
|---|---|---|---|
| *Car* | NM_009803 | GCTGCAAGGGCTTCTTCAGA | CCTTCCAGCAAACGGACAGA |
| *Cyp2b10* | NM_009999 | TTTCTGCCCTTCTCAACAGGAA | ATGGACGTGAAGAAAAGGAACAAC |
| *Cyp2c55* | NM_028089 | TTGTGGAAGAGCTAAGAAAAGCAAAT | GAGCACAGCTCAGGATGAATGT |
| *Fmo3* | NM_008030 | AAGAAAGGAAGACAAAGAAAAGGCA | AGCTCCAATGATGGCCACTT |

**Additional File 2: $^{1}H$ and partial $^{13}C$ assignments for identified metabolites in plasma samples.** Signals highlighted in bold were used for determination of area under the curves. Keys: s: singlet; d: doublet; t: triplet; q: quadruplet; m: multiplet; dd: doublet of doublet.

*: tentative assignment

| Metabolite | Chemical function | δ $^{1}H$ (ppm) | Multiplicity | δ $^{13}C$ (ppm) |
|---|---|---|---|---|
| Cholesterol (mainly in HDL) | | **0.66** | **s (broad)** | |
| lipid in LDL | ($CH_3(CH_2)_n$) | **0.85** | **s (broad)** | 24.6 |
| | (CH2)n | 1.23 | s (broad) | 32.0 |
| lipid in VLDL | ($CH_3CH_3CH_2C$=) | **0.87** | **t** | 16.5 |
| | $CH_2CH_2CH_2CO$ | 1.29 | m | |
| | $CH_2CH_2CO$ | 1.59 | m | 27.0 |
| Leucine | | 0.95 | t | |
| Valine | | 0.99 | d | |
| | | 1.05 | d | 20.3 |
| Isoleucine | | 1.01 | d | |
| Ethanol | | 1.19 | t | 19.4 |
| | | 3.65 | q | 60.4 |
| lactate | | **1.33** | **d** | 22.4 |
| | | 4.12 | q | 71.3 |
| alanine | | 1.48 | d | 19.0 |
| unknown | | 1.73 | m | 28.3 |
| acetate | | 1.92 | s | 26.0 |
| lipid | $CH_2C$=C | 2.0 | m | 29.0 |
| | $CH_2C$=C | 2.04 | m | |
| methionine | | 2.13 | s | 17.1 |
| lipid | $CH_2CO$ | 2.24 | t | 35.7 |
| 3-hydroxybutyrate | | 2.31 | dd | |
| succinate* | | 2.4 | s | |
| glutamine | | 2.45 | m | 33.5 |
| unknown | | 2.53 | s | 34.7 |
| methylamine | | 2.55 | s | 24.7 |
| unknown | | 2.69 | s | 28.5 |
| dimethylamine* | | 2.71 | s | |
| lipid | | 2.76 | m | 28.3 |
| creatine | | 3.04 | s | 42.2 |
| glycerophosphocholine | | 3.22 | s | 56.7 |
| β-glucose | | 3.25 | m | 77.2 |
| | | 3.47 | m | 79.0 |
| | | 3.9 | m | 63.5 |
| | | 4.65 | d | 98.9 |
| α-glucose | | 3.41 | m | 72.5 |
| | | 3.53 | m | 72.0 |
| | | 3.71 | m | 63.0 |
| | | 3.78 | m | 63.7 |
| | | 3.85 | m | 74.3 |

| | | | |
|---|---|---|---|
| | **5.25** | **d** | 95.4 |
| unsaturated lipids | 5.3 | m | 132.2 |
| fumarate* | 6.52 | s | |
| tyrosine | 6.9 | d | 118.7 |
| | 7.19 | d | 134.7 |
| unknown | 7.05 | s | |
| phenylalanine | 7.34 | m | 146.2 |
| | 7.43 | t | 146.4 |
| unknown | 7.75 | s | 147.0 |

**Additional File 3: $^{1}H$ and partial $^{13}C$ assignments for identified metabolites in liver samples.** Signals highlighted in bold were used for determination of area under the curves. Keys: s: singlet; d: doublet; t: triplet; q: quadruplet; m: multiplet; dd: doublet of doublet.

x*: tentative assignment

| Metabolite | δ $^{1}H$ (ppm) | Multiplicity | δ $^{13}C$ (ppm) |
|---|---|---|---|
| Bile acids (mixed) | 0.6-0.75 | s | |
| | 0.92 | s | |
| Bile acids (tauroconjugated) | 0.72 | s | |
| | 0.92 | s | |
| | 3.08 | t | |
| Leucine | 0.96 | t | 23.6 |
| | 1.72 | m | |
| | 3.74 | m | |
| Valine | 0.99 | d | 19.5 |
| | 1.05 | d | 20.8 |
| | 2.29 | | |
| | 3.62 | | |
| Isoleucine | 0.94 | t | |
| | 1.01 | d | |
| | 1.26 | m | |
| | 1.48 | m | |
| | 2.0 | m | |
| | 3.65 | d | |
| 3-hydroxybutyrate | 1.19 | d | 24.4 |
| | 2.32 | dd | |
| | 2.4 | dd | |
| | 4.14 | m | |
| Lactate | 1.33 | d | 22.8 |
| | 4.11 | q | |
| Threonine | 1.33 | d | 22.3 |
| | 3.61 | | |
| | 4.23 | | |
| Alanine | 1.49 | d | 19.1 |
| | 3.79 | | |
| Ornithine | 1.72 | m | |
| | 1.93 | m | |
| | 3.03 | t | 41.9 |
| | 3.77 | t | |
| Acetate | 1.93 | s | |
| L-glutamate | 2.06 | m | 29.8 |
| | 2.35 | m | 36.4 |
| | 3.76 | dd | |
| L-glutamine | 2.13 | m | 29.3 |
| | 2.45 | m | |
| | 3.77 | t | |
| Glutathion (oxidized) | 2.17 | t | 29.1 |
| | **2.53** | **m** | 34.2 |

| | | | |
|---|---|---|---|
| | 2.98 | dd | |
| | 3.31 | m | 41.6 |
| | 3.76 | m | |
| | 4.75 | m | |
| Glutathion (reduced) | 2.17 | m | |
| | **2.56** | **m** | |
| | 2.95 | m | |
| | 3.76 | m | |
| | 4.56 | dd | |
| Succinate | 2.42 | s | 36.8 |
| Hypotaurine | **2.65** | **t** | |
| | 3.36 | t | |
| L-aspartic acid | 2.66 | dd | |
| | 2.8 | dd | |
| | 3.89 | dd | |
| Dimethylamine | 2.72 | s | 37.6 |
| Trimethylamine | **2.89** | **s** | |
| Dimethylglycine | 2.93 | s | 46.5 |
| Creatine | 3.04 | s | 39.9 |
| | 3.94 | s | 56.6 |
| Choline | 3.2 | s | 56.7 |
| | 3.55 | | |
| O-phosphocholine | 3.21 | s | 56.6 |
| | 3.58 | m | |
| | 4.13 | m | |
| Betaine | 3.27 | s | 55.9 |
| | 3.88 | s | |
| Taurine | 3.28 | t | 50.4 |
| | 3.44 | t | 38.5 |
| Methanol | 3.36 | s | 51.7 |
| β-glucose | 3.51 | m | |
| | 3.75 | | |
| | 4.66 | d | 98.7 |
| AMP | 4.01 | dd | |
| | 4.36 | dd | |
| | 4.5 | dd | |
| | 6.14 | d | 89.6 |
| | 8.27 | s | |
| | 8.62 | s | |
| α-glucose | 3.45 | m | |
| | 3.56 | | |
| | 3.72 | | |
| | 3.84 | | |
| | 3.97 | | |
| | 5.25 | d | 94.8 |
| Glycine | 3.56 | s | 44.4 |
| UDP-glucose | 5.6 | dd | |
| | 5.95 | d | |
| | 7.93 | d | |

| | | | |
|---|---|---|---|
| UDP-glucuronate | 5.6 | dd | |
| | 5.98 | d | |
| | 7.96 | d | |
| Uridine | 5.89 | d | |
| | 5.9 | d | |
| | 7.89 | d | |
| NADP+ | 6.03 | s | |
| | 6.1 | d | |
| | 8.15 | d | |
| | 8.42 | s | |
| | 8.584 | s | |
| | 8.82 | s | |
| | 9.12 | d | |
| | 9.3 | s | |
| NAD+ | 6.04 | d | 89.5 |
| | 6.09 | d | 102.6 |
| | 8.18 | s | |
| | 8.2 | dd | |
| | 8.44 | s | |
| | 8.84 | d | |
| | 9.15 | d | |
| | 9.34 | s | |
| Inosine | 6.11 | d | 91.1 |
| | 8.23 | s | |
| | 8.34 | s | |
| Fumarate | 6.52 | s | |
| Tyrosine | 6.87 | d | |
| | 7.19 | | 132.5 |
| Phenylalanine | 3.12 | dd | |
| | 3.26 | dd | |
| | 7.33 | m | |
| | 7.38 | m | |
| | 7.43 | m | |
| Nicotinurate | 3.99 | s | |
| | 7.6 | dd | |
| | 8.25 | m | |
| | 8.72 | dd | |
| | 8.94 | s | |
| Nicotinamide | 7.61 | dd | |
| | 8.23 | dd | |
| | 8.7 | dd | |
| | 8.95 | s | |

**Additional File 4: Impact of CAR activation on hepatic transcripts in *Car$^{+/+}$* *vs.* *Car$^{-/-}$* mice.** A) PCA score plots of the whole liver transcriptomic dataset in *Car$^{+/+}$* and *Car$^{-/-}$* mice (n=6/group). B) Venn diagram representing the number of genes significantly modulated by TCPOBOP in the liver of *Car$^{-/-}$* *vs.* *Car$^{+/+}$* male and female mice.

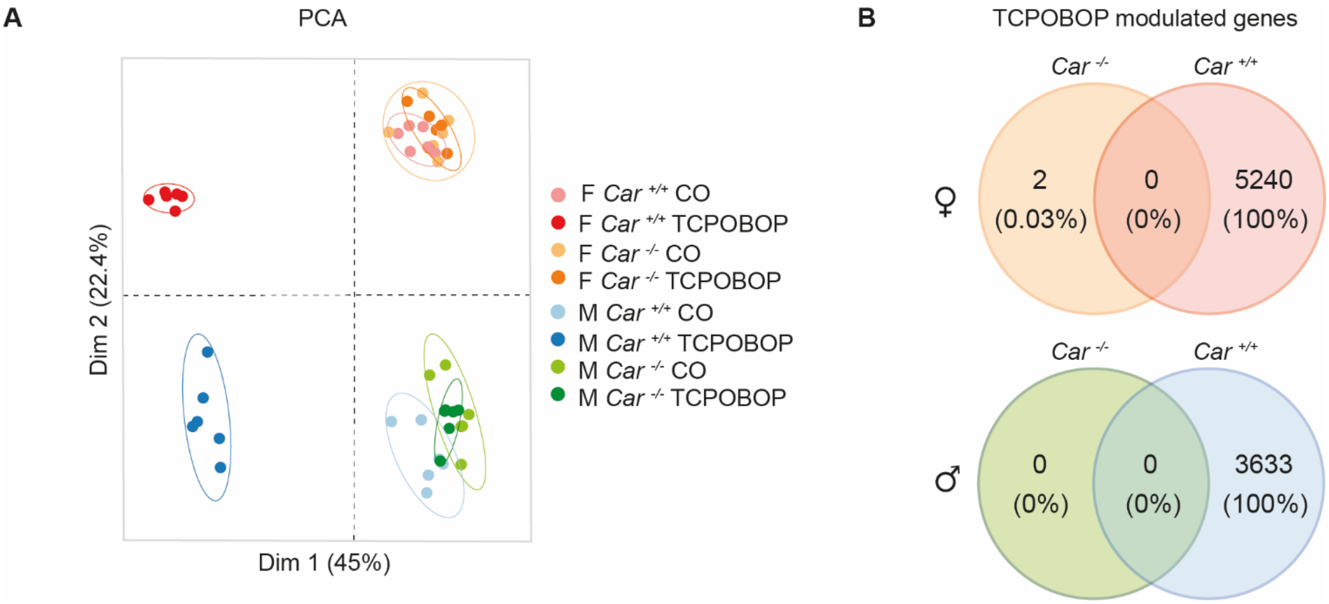

**Additional File 5: Majority of TCPOBOP-modulated genes are common between sexes.** A) Venn diagram representing the number of genes significantly modulated by TCPOBOP in *Car* $^{+/+}$ male and female mice liver considering significant regulation at $P_{adj}$<0.05 and 3 different fold-change thresholds (TCPOBOP *vs.* CO). B) Venn diagram representing the number of genes significantly modulated by TCPOBOP in *Car*$^{+/+}$ male and female mouse liver considering significant regulation at $P_{adj}$ <0.01 and 3 different fold-change thresholds (TCPOBOP *vs.* CO).

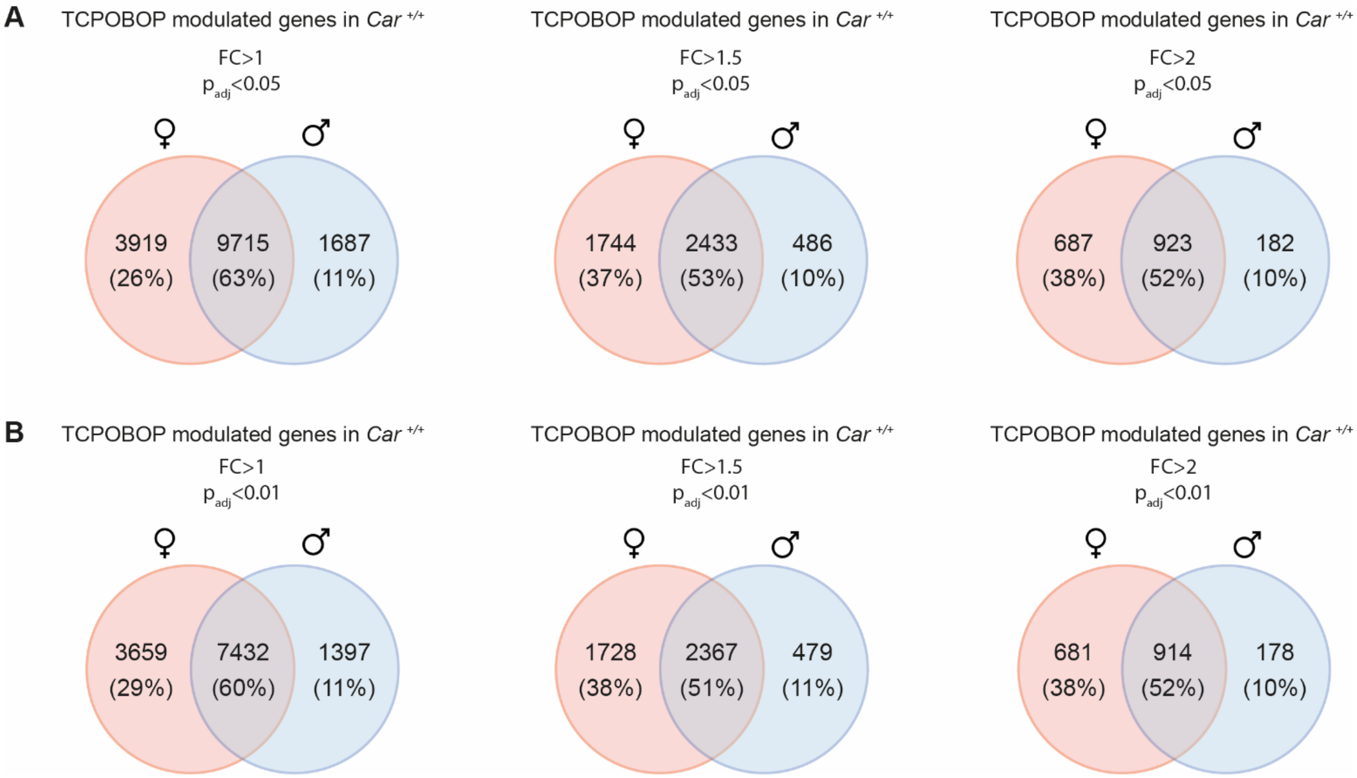

**Additional File 6: Fold-changes, p-values and pathway enrichment analysis for all TCPOBOP modulated genes (related to Fig. 2c).** A) List of genes contained in cluster 1 with their Log(fold-change) values and adjusted p-values in *Car*$^{+/+}$ males and females. B) Gene Ontology (GO) term enrichment analysis using the list of genes contained in cluster 1. C) List of genes contained in cluster 2 with their Log(fold-change) values and adjusted p-values in males and females *Car*$^{+/+}$. D) GO term enrichment analysis using the list of genes contained in cluster 2. Metabolic pathways were considered as significantly enriched when $P_{adj} < 0.05$.

**Additional File 7: Expression of previously-described CAR target genes in both sexes (related to Fig. 2e-g).** Table presenting list of genes well-described in the literature as CAR-sensitive genes with their Log(fold-change) values and adjusted p-values in males and females *Car*$^{+/+}$. Data were derived from liver microarray data.

**Additional File 8: Fold-changes, p-values and pathway enrichment analysis for genes modulated by TCPOBOP in a sex-dependent manner (related to Fig. 3b).** A) Genes contained in cluster 1 with their Log(fold-change) values and adjusted p-values in *Car*$^{+/+}$ males and females. B) GO term enrichment analysis C) Genes contained in cluster 2 with their Log(fold-change) values and adjusted p-values in males and females *Car*$^{+/+}$ groups. D) GO term enrichment analysis using the list of genes contained in cluster 2. E) List of genes contained in cluster 3 with their Log(fold-change) values and adjusted p-values in males and females *Car*$^{+/+}$ groups. F) GO term enrichment analysis. G) List of genes contained in cluster 4 with their Log(fold-change) values and adjusted p-values in males and females *Car*$^{+/+}$ groups. H) GO term enrichment analysis.

**Additional File 9: $^{1}$H-NMR-based metabolomics in plasma samples from *Car*$^{-/-}$ mice.** A) Coefficient plots related to the PLS-DA models discriminating between $^{1}$H-NMR-based plasma spectra from TCPOBOP *vs.* CO *Car*$^{-/-}$ males. Parameters of the PLS-DA model: $Q^2Y=0.1$, $p>0.05$. B) Coefficient plots related to the PLS-DA models discriminating between $^{1}$H-NMR-based plasma spectra from TCPOBOP *vs.* CO *Car*$^{-/-}$ females. Parameters of the PLS-DA model: $Q^2Y=-0.4$, $p>0.05$.

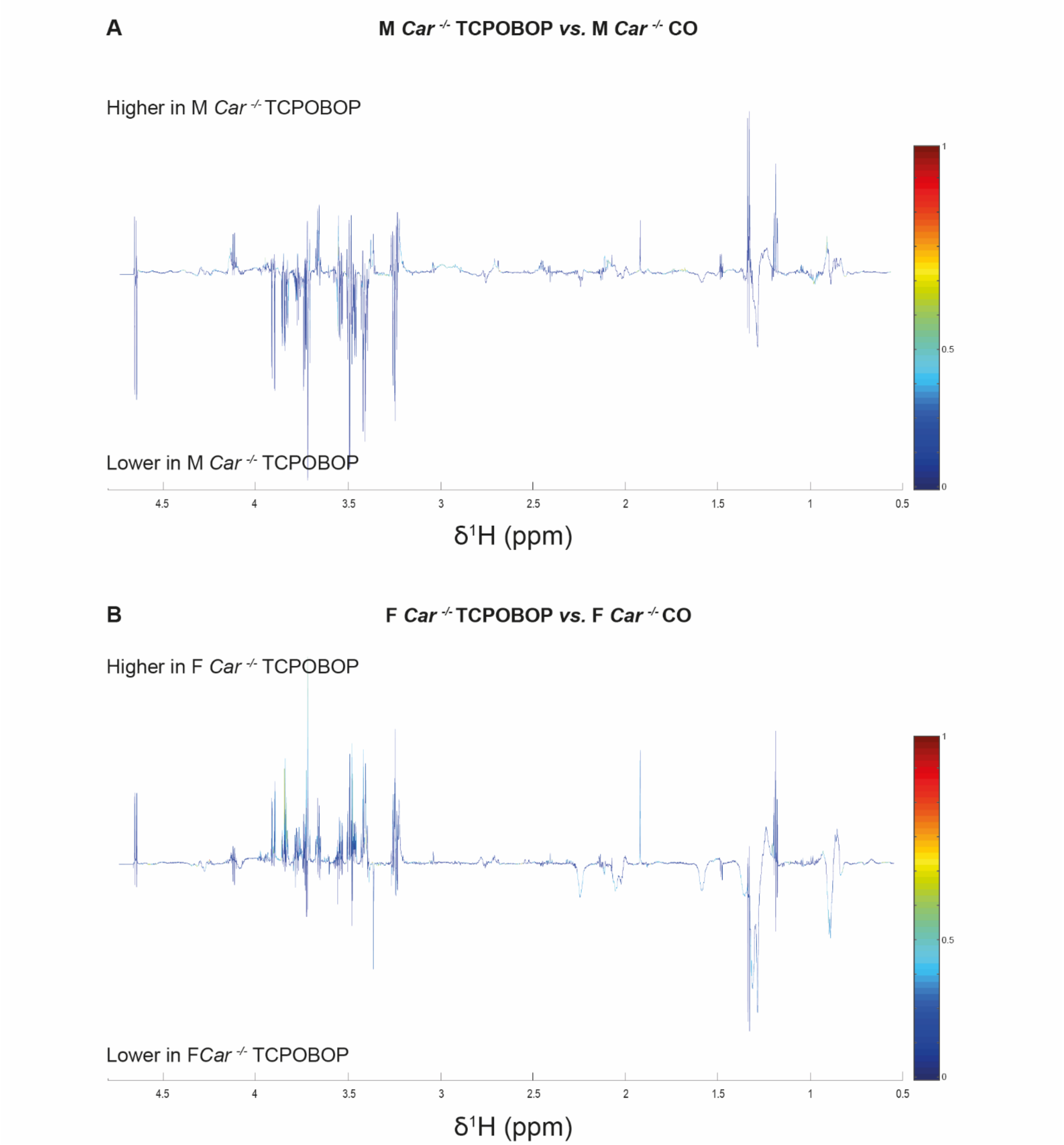

**Additional File 10: Effect of CAR activation on plasma cholesterol, LDL-cholesterol and HDL-cholesterol.** A) Plasma cholesterol concentration was determined on plasma collected at the end of the experiment. B) Plasma concentration of low-density-lipoprotein (LDL) was determined on plasma collected at the end of the experiment. C) Plasma concentration of high-density-lipoprotein (HDL) was determined on plasma collected at the end of the experiment. Results are given as the mean ± SEM. *treatment effect, #sex effect. * or # p<0.05, ** or ## p<0.01, *** or ### p<0.001.

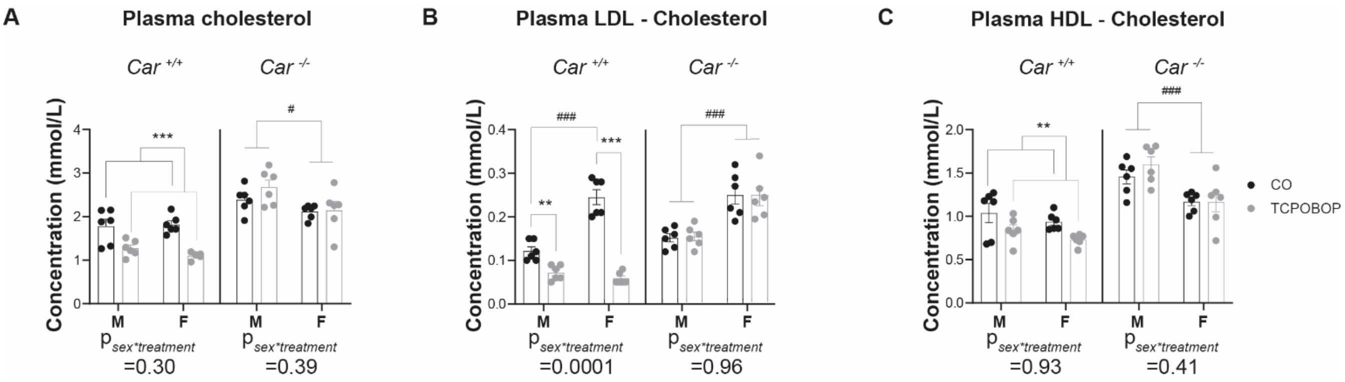

**Additional File 11: Effect of CAR activation on hepatic mRNA expression of genes involved in lipoprotein metabolism in *Car*$^{-/-}$ mice.** A) Fold-change (TCPOBOP- *vs.* CO-treated *Car*$^{-/-}$ mice) of hepatic expression for genes involved in lipoprotein metabolism (derived from microarray results).

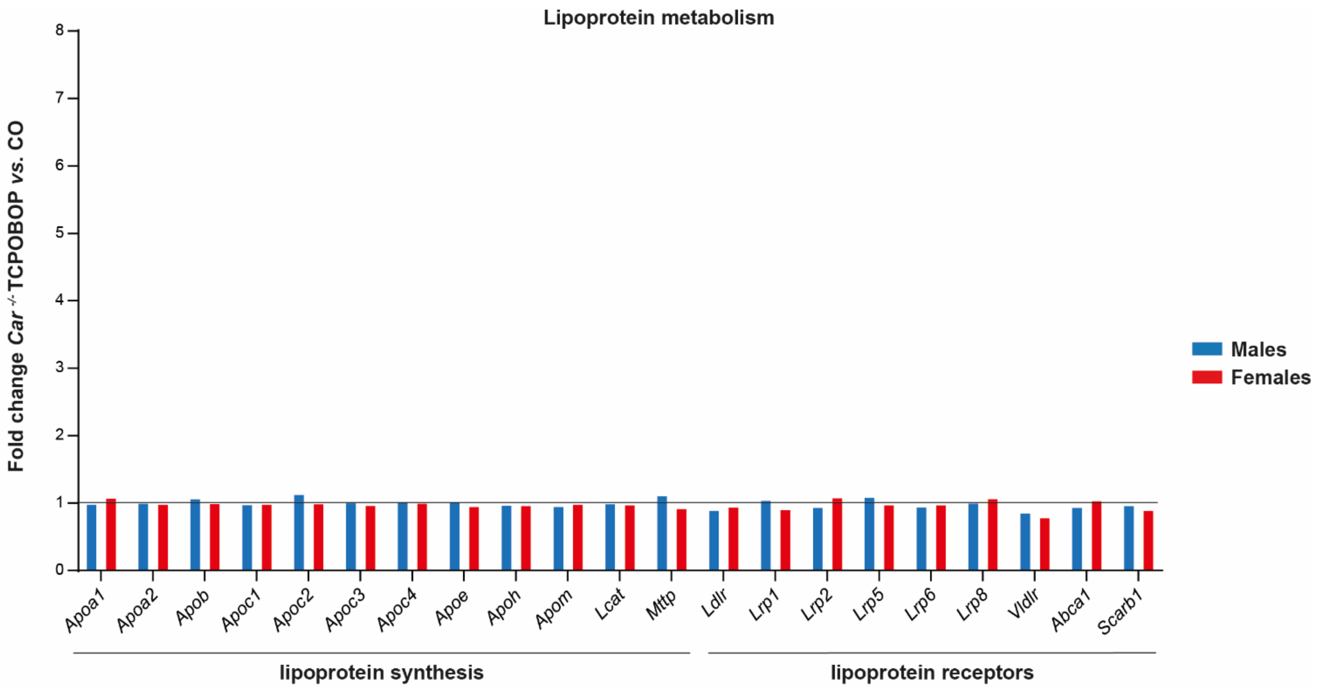

**Additional File 12: Pharmacological activation of CAR modulates expression of genes involved in bile acid metabolism and cholesterol metabolism.** A) Fold-change (TCPOBOP- *vs.* CO-treated *Car*$^{+/+}$ mice) of hepatic expression for genes involved in bile acid metabolism (derived from microarray results). B) Fold-change (TCPOBOP- *vs.* CO-treated *Car*$^{+/+}$ mice) of hepatic expression for genes involved in cholesterol metabolism (derived from microarray results). C) Fold-change (TCPOBOP- *vs.* CO-treated *Car*$^{-/-}$ mice) of hepatic expression for genes involved in bile acid metabolism (derived from microarray results). D) Fold-change (TCPOBOP- *vs.* CO-treated *Car*$^{-/-}$ mice) of hepatic expression for genes involved in cholesterol metabolism (derived from microarray results). *treatment effect. * $p_{adj}<0.05$, ** $p_{adj}<0.01$, *** $p_{adj}<0.001$.

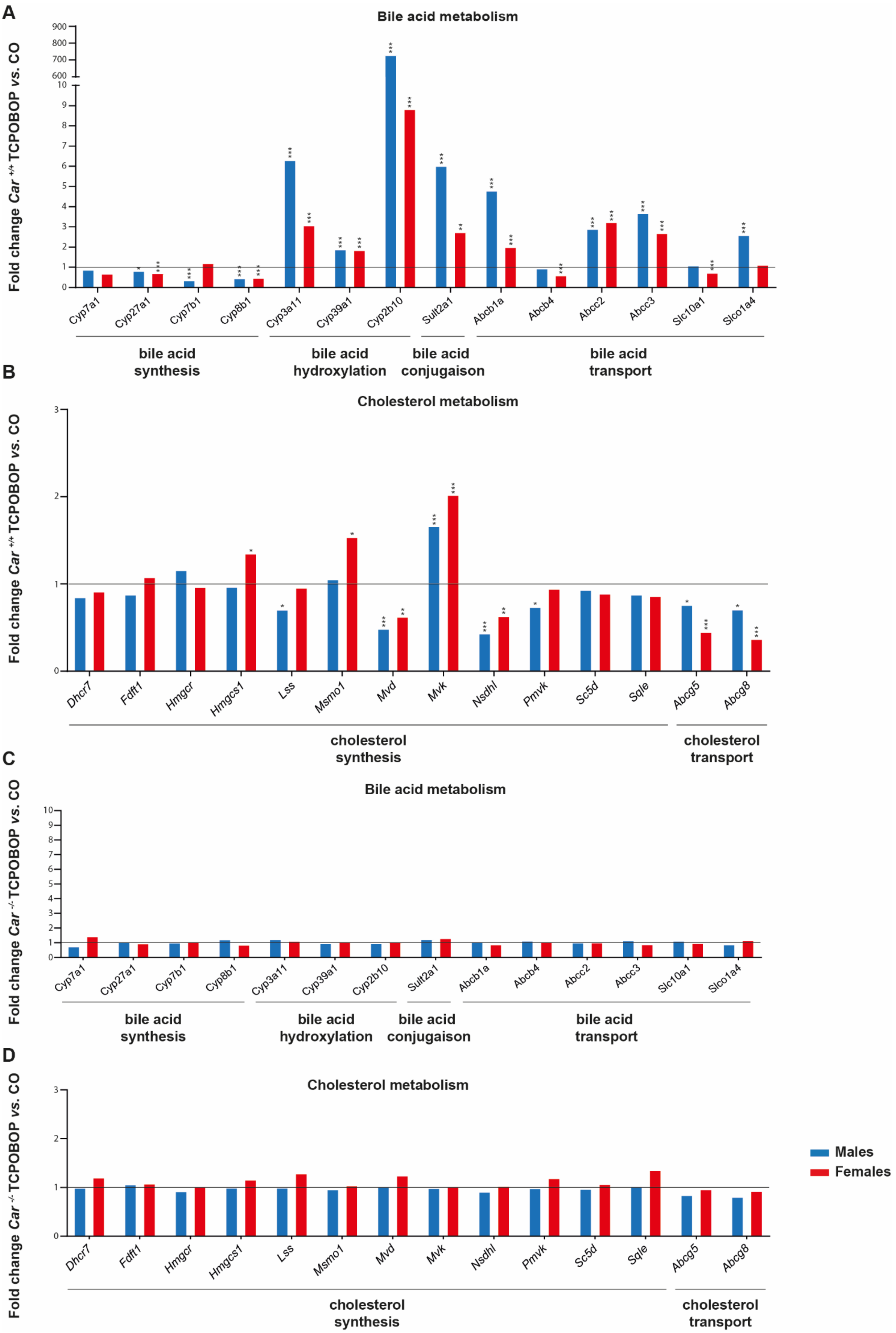
A
Bile acid metabolism
Fold change Car +/+ TCPOBOP vs. CO
Cyp7a1
Cyp27a1
Cyp7b1
Cyp8b1
Cyp3a11
Cyp39a1
Cyp2b10
Sult2a1
Abcb1a
Abcb4
Abcc2
Abcc3
Slc10a1
Slco1a4
bile acid synthesis
bile acid hydroxylation
bile acid conjugaison
bile acid transport
B
Cholesterol metabolism
Fold change Car +/+ TCPOBOP vs. CO
Dhcr7
Fdft1
Hmgcr
Hmgcs1
Lss
Msmo1
Mvd
Mvk
Nsdhl
Pmvk
Sc5d
Sqle
Abcg5
Abcg8
cholesterol synthesis
cholesterol transport
C
Bile acid metabolism
Fold change Car -/- TCPOBOP vs. CO
Cyp7a1
Cyp27a1
Cyp7b1
Cyp8b1
Cyp3a11
Cyp39a1
Cyp2b10
Sult2a1
Abcb1a
Abcb4
Abcc2
Abcc3
Slc10a1
Slco1a4
bile acid synthesis
bile acid hydroxylation
bile acid conjugaison
bile acid transport
D
Cholesterol metabolism
Fold change Car -/- TCPOBOP vs. CO
Dhcr7
Fdft1
Hmgcr
Hmgcs1
Lss
Msmo1
Mvd
Mvk
Nsdhl
Pmvk
Sc5d
Sqle
Abcg5
Abcg8
cholesterol synthesis
cholesterol transport
Males
Females

**Additional File 13: $^{1}$H-NMR-based metabolomics in liver samples from *Car*$^{-/-}$ mice.** A) Coefficient plots related to the PLS-DA models discriminating between $^{1}$H-NMR-based liver spectra from TCPOBOP *vs.* CO *Car*$^{-/-}$ males. Parameters of the PLS-DA model: $Q^2Y=-0.18$, $p>0.05$. B) Coefficient plots related to the PLS-DA models discriminating between $^{1}$H-NMR-based liver spectra from TCPOBOP *vs.* CO *Car*$^{-/-}$ females. Parameters of the PLS-DA model: $Q^2Y=-0.28$, $p>0.05$.

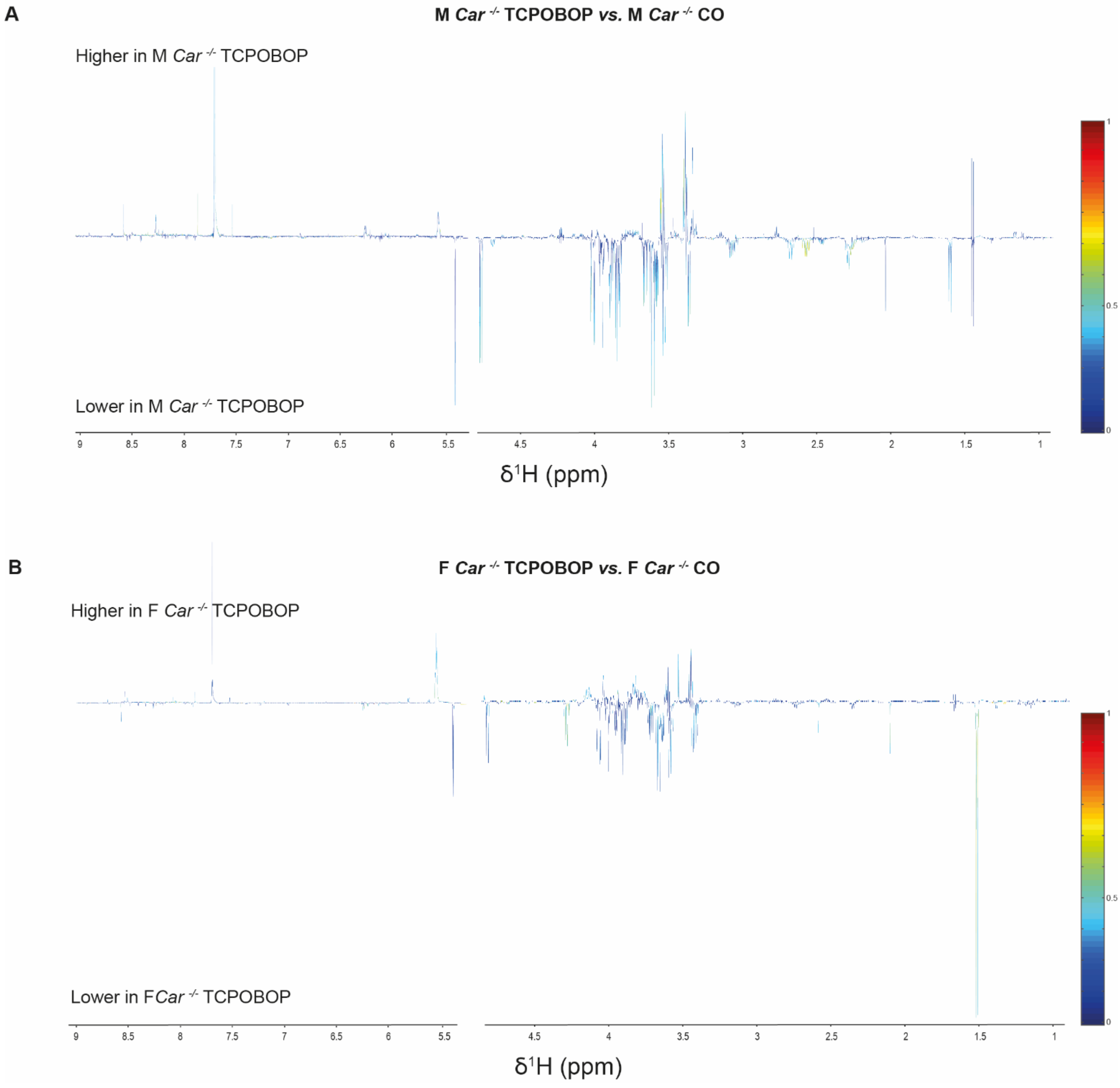
